# Phospholipases D1 and D2 regulate cell cycling in primary prostate cancer cells and are differentially associated with the nuclear matrix

**DOI:** 10.1101/2021.05.19.444795

**Authors:** Amanda R Noble, Karen Hogg, Sylvain Bourgoin, Dawn Coverley, Leanne Archer, Fiona Frame, Norman J Maitland, Martin G Rumsby

## Abstract

Phospholipases D1 and D2 (PLD1/2) have been implicated in tumorigenesis. We previously detected higher expression of PLD in the nuclei of patient-derived prostate cancer (PCa) cells and prostate cancer cell lines. Here we have examined whether PLD1 or PLD2 are associated with the nuclear matrix and influence cell cycling. PLD1/PLD2 were detected by qualitative immunofluorescence in cultured PCa cells and extracted with a standardised protocol to reveal nuclear matrix-associated proteins. The effects of isoform-specific inhibition of PLD1or PLD2 on PCa cell cycle progression were analysed by flow cytometry. PLD2 mainly co-localised with the nucleolar marker fibrillarin in PCa cells. However, even after complete extraction, some PLD2 remained associated with the nuclear matrix. Inhibiting PLD2 effectively reduced PCa cell cycling into and through S phase. In contrast, PLD1 inhibition effects were weaker, and a subpopulation of cycling patient-derived PCa cells was unaffected by PLD1 inhibition. When associated with the nuclear matrix PLD2 could generate phosphatidic acid to regulate nuclear mTOR and control downstream transcriptional events. The association of PLD2 with the nucleolus also implies a role in stress regulation. The cell cycling results highlight the importance of PLD2 inhibition as a novel potential prostate cancer therapeutic mechanism by differential regulation of cell proliferation.

## 1. Introduction

The two main isoforms of mammalian phospholipase D (PLD), PLD1 and PLD2, hydrolyse chiefly phosphatidylcholine (PtdCho) to a base and the signalling lipid phosphatidic acid (PtdOH). Both PLD1 and PLD2 are implicated in tumorigenesis since total PLD activity and PLD1/PLD2 protein expression are elevated in many cancers and often correlate with prognosis^1-8^. In prostate cancer (PCa) we have reported that PLD2 protein expression correlates with increasing Gleason score (GS) from GS6 to GS8 but is decreased in GS9 tissue where gland structure is not evident^9^. Stable fibroblast cell lines over-expressing PLD1 or PLD2 show increased proliferation, enhanced colony formation in soft agar and induce undifferentiated sarcomas in nude mice^10^. Survival and migration signals in human breast cancer cells and androgen-insensitive PCa cell lines are both associated with higher PLD activity^11,12^. In bone metastases-derived PC3 and C4-2B prostate cell lines, increased PLD activity and PLD1/PLD2 expression contributes to tumorigenesis^13^. PLD2 also regulates exosome secretion by C4-2B and PC3 cells resulting in increased osteoblast activity^14^.

The involvement of PtdOH in the recruitment and activation of mTOR (mechanistic [mammalian] target of rapamycin)^15^, has defined a mechanistic role for PLD1 and PLD2 in tumorigenesis^16,17^. Active mTOR in the cytosol exists as complexes; mTORC1 contains Raptor and mLST8, and mTORC2 contains Rictor, mLST8 and sin1.^18,19^ PLD1 is detected at a perinuclear site in cells and migrates to the plasma membrane on activation^20,21^. In contrast, PLD2 is located at the plasma membrane and is associated in lipid rafts promoting receptor endocytosis^22^,^23^. In recent work we also detected PLD1 and PLD2 protein expression in the nuclei^9,24^ of patient-derived PCa cells and PCa cell lines in agreement with occasional reports in other cell types^4,25-27^.

A role for both PLD1 and PLD2 in cell cycle control has been defined. For example, fibroblasts overexpressing PLD1 or PLD2 show an increased proportion of cells in S phase while the level of cyclin D3 protein, an activator of the G1 to S phase transition, is also elevated^10^. Increased expression of either PLD1 or PLD2 prevents cell cycle arrest by high intensity Raf signals^28^ while PLD1 and PLD2 repress p21 gene transcription stimulating cell growth, and resulting in tumorigenesis^29^. In colorectal cancer cells inhibition of PLD induces cell cycle arrest^30^. These effects of PLD on the cell cycle occur through mTOR, which operates checkpoints in cell cycle progression and requires PtdOH for activity^15,17,31^. mTOR is also present in the nucleus (nmTOR), bound to the promoters of RNA polymerase I and III genes where it regulates transcription^32^. In skeletal muscle, nmTOR binds to the promoters of some mitochondrial genes^33^. In PCa, nmTOR functions as a transcriptional integrator of androgen signalling pathways and increased nmTOR correlates with a poor prognosis^34^.

To confirm and extend our findings on the role of PLD1 and PLD2 in PCa cell nuclei we have now examined the association of PLD1 and PLD2 with the nucleus by treating patient-derived PCa cells and PC3 cells in a well-documented fractionation procedure which exposes core nuclear matrix (NM) proteins^35-37^ Nuclei treated by this fractionation scheme leave a protein-rich matrix which includes actin and fibrillarin^38^ and nuclear lamina (NL) proteins ^39^. We have begun to examine what role PLD1 and PLD2 might play in the nucleus by examining the effect their inhibition has on prostate PCa cell cycling.

## 2. Materials and Methods

### 2.1. Cell isolation and culture

The PC3 prostate epithelial cell line was cultured as described in Noble et al^9,24^. Primary PCa epithelial cells were purified from human prostate tissue samples which were obtained with patient consent and full ethical approval (South Yorkshire Research Ethics Committee, Yorkshire and the Humber, REC:07/H1304/121) as previously stated^24^. The cells were grown on collagen 1-coated 10cm dishes in keratinocyte serum-free medium with supplements of L– glutamine, bovine pituitary extract, epidermal growth factor, stem cell factor, cholera toxin, leukaemia inhibitory factor and granulocyte-macrophage colony-stimulating factor at 37 °C with 5% CO_2_. Cells were initially co-cultured with irradiated (60Gy) mouse embryonic fibroblasts (STO). Subsequent passages were free of STOs and all cultures were used at the lowest practical passage number (p2-p5).

### 2.2. Association of PLD1 and PLD2 with the nuclear matrix

Patient-derived primary PCa cells and PC3 cells were cultured on collagen-coated chamber slides as above, and were then subjected to a standardised detergent, high salt, DNase and RNase extraction procedure^35-37^ to expose the nuclear matrix (Fig. 1). After each stage, cells were examined for PLD1 and PLD2 by qualitative immunofluorescence. DAPI was used to define nuclear DNA, lamin the NL and fibrillarin the nucleolus. All images were captured under identical conditions to allow comparisons to be made through the extraction series.

**Figure 1.**
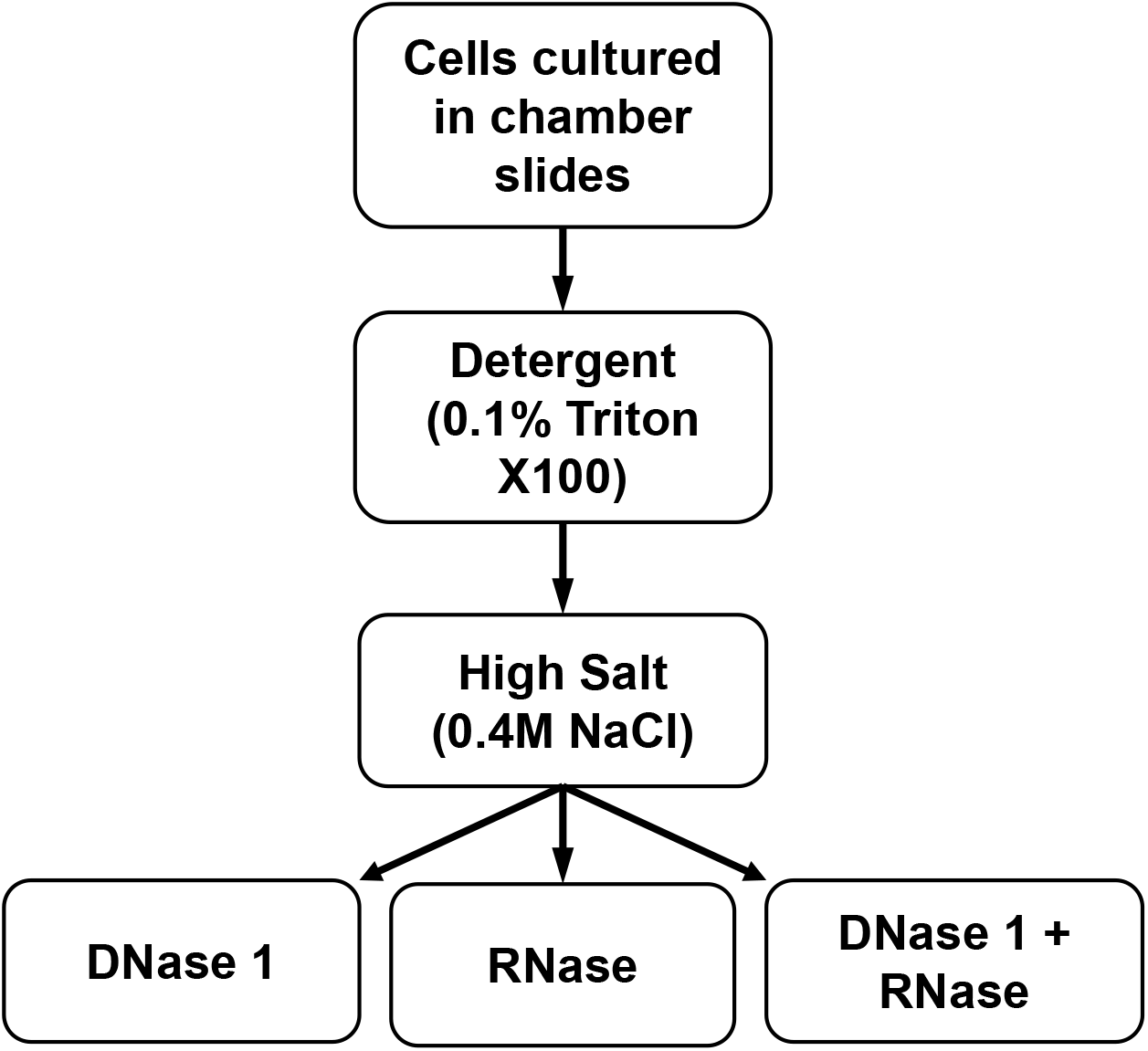
Nuclear Matrix Extraction procedure^35-37^. Cells cultured in chamber slides were treated with detergent revealing detergent-resistant PLD1/2, followed by 0.4M NaCl revealing high salt-resistant PLD1/2 then either DNAse 1, RNase or DNAse 1 + RNase revealing nuclease-resistant proteins.

### 2.3. Immunofluorescence

Intact PCa cells were fixed in 4% paraformaldehyde, rinsed with PBS, permeabilised with 0.5% Triton X-100 and rinsed again. Untreated or extracted cells from 2.2 above were rinsed, blocked in 10% goat serum in PBS for 60 minutes and then treated with primary antibody in 10% goat serum overnight at 4°C. Next day the cells or cell residues were rinsed and the appropriate Alexafluor secondary antibody added for 1 hour at room temperature, followed by rinses. The chambers were removed and the slides were mounted using Vectashield with DAPI (Vector laboratories, Peterborough, UK) and examined using a Nikon Eclipse TE300 fluorescence microscope (Nikon, Surrey, UK). Primary antibodies were a commercial anti-PLD1 antibody (Santa Cruz, sc25512, validated by Bruntz et al^23^ and Scott et al^40^), a polyclonal anti-PLD2 antibody (PLD2-26, validated by Denmat-Ouisse et al^41^), a mouse anti-fibrillarin antibody (Abcam ab4566), an anti-lamin B2 antibody (Invitrogen 33-2100), a rabbit polyclonal anti-Raptor antibody (Proteintech, 20984-1-AP), a rabbit polyclonal anti-Rictor antibody (Proteintech, 27248-1-AP) and a mouse monoclonal anti-mTOR antibody (Proteintech, 66888-1-Ig) all used at 1: 100. Secondary antibodies were a goat anti-rabbit Alexa Fluor 568 (A11036, Thermofisher) and a goat anti-mouse Alexa Fluor 488 (A11029, Thermofisher).

### 2.4. PLD inhibition and the cell cycle

The effects of PLD inhibition on cell cycle progression were measured by flow cytometry using the Click-iT Plus EdU Pacific Blue Flow Cytometry Kit (ThermoFisher Scientific C10636) according to the manufacturer’s instructions. PC3 cells (6×10^4^) were plated in 12 well plates with the appropriate medium. The following day the cells were treated with vehicle (DMSO), 10μM PLD1 inhibitor (EVJ), 10μM PLD2 inhibitor (JWJ) or both. At 24 and 48 hour time points 5-ethynyl-2’-deoxyuridine (EdU) was added to separate samples. 4 hours later cells were harvested according to the manufacturer’s protocol. Primary cells purified from three different PCa biopsy samples were plated at 6×10^4^ cells/well in 6 well collagen-coated plates with stem cell media; 4×10^4^ cells/well were plated for the control wells. The following day the cells were treated with vehicle (DMSO), PLD 1 inhibitor EVJ, PLD2 inhibitor JWJ at their IC_50_ values, ie EVJ 11.4 μM, JWJ 6.4 μM, and in combination at half their IC_50_ values (EVJ 5.7μM, JWJ 3.2μM), these concentrations being based on results from our previous inhibition studies.^9,24^ After 48 hours 5-ethynyl-2’-deoxyuridine (EdU) was added and after a further 24 hours the cells were harvested according to the manufacturer’s protocol. Results were acquired on a CyAn ADP flow cytometer using Summit software. EdU-Pacific blue was excited by 405nm laser-emitted photons detected at 450/50nm bandwidth: DNA-propidium iodide (PI) was excited by 488nm laser and emitted photons detected at 613/20 bandwidth. The cell population of interest was gated using scatter plots FS Lin / SS Log and FS Lin / Pulse Width. PI Lin Area / EdU Log was used to determine the percentage of EdU^+^-cells (ie cells cycling into or passing through S phase), cells in G1+G2/M phases of the cell cycle, and, where identified, a subG1 population of dying cells with fragmented DNA. The PLD1 inhibitor EVJ (VU0359595) and the PLD2 inhibitor JWJ (VU0364739) were gifts from the late Professor Alex Brown, Vanderbilt University, USA^42-44^.

## 3. Results

### 3.1. PLD1 and PLD2 are associated with the nucleus in PC3 cells

In untreated PC3 cells (Fig. 2A, a), PLD1 protein is present in the cytosol (white arrow) and both diffusely throughout the nucleus and as a concentrated spot (blue arrows). With DAPI to define chromatin, a DAPI/PLD1 overlay (Fig. 2A, c) shows this cytosolic/nuclear distribution of PLD1 more clearly. Detergent, high salt and DNase treatment (Fig. 2A, d) exposes PLD1 in both the cytosol and nuclei so that fluorescence is enhanced; this is especially true for the concentrated spot of PLD1 in nuclei (Fig. 2A, d, blue arrows). PLD1 protein detection in the cytosol is also enhanced (Fig. 2A, d, white arrows). Staining for the intermediate filament protein lamin defines the NL (Fig. 2A, e). A lamin/PLD1 overlay (Fig. 2A, f) confirms that the concentrated spot of PLD1 observed in Fig. 2A, d, is within the nucleus. Nuclear PLD1 was not apparent in PC3 cells after subsequent RNase treatment though some cytoplasmic PLD1 was still faintly detected (Fig. 2A, g, i, white arrows). Results for each stage of the extraction sequence for PLD1 in PC3 cells are shown in Supplementary Fig. 1. A different expression pattern is noted for PLD2 protein, which is detectable in both the nuclei (blue arrows) and cytosol (white arrows) of untreated PC3 cells (Fig. 2B, a). Chromatin is defined by DAPI (blue, Fig. 2B, b) and nucleoli by fibrillarin (green, Fig. 2B, c); several nuclei have more than one nucleolus (green arrows). In an overlay with DAPI (d) cytosolic PLD2 (white arrows) and nucleoli (green arrows) are still apparent. After detergent and high salt treatment, cytosolic PLD2 protein is concentrated in a mainly perinuclear region (e, white arrow), whereas in nuclei, the protein is diffusely distributed and as single or several concentrated spots (e, blue arrows) which match the positions of nucleoli as defined by staining for fibrillarin (g, green arrows). In an overlay (h) with DAPI (blue) to define chromatin (f) a rim of cytosolic PLD2 (white arrows) remains while the concentrated spots of nuclear PLD2 appear yellowish indicating co-localisation with the nucleolus marker fibrillarin (blue arrows).

**Figure 2.**
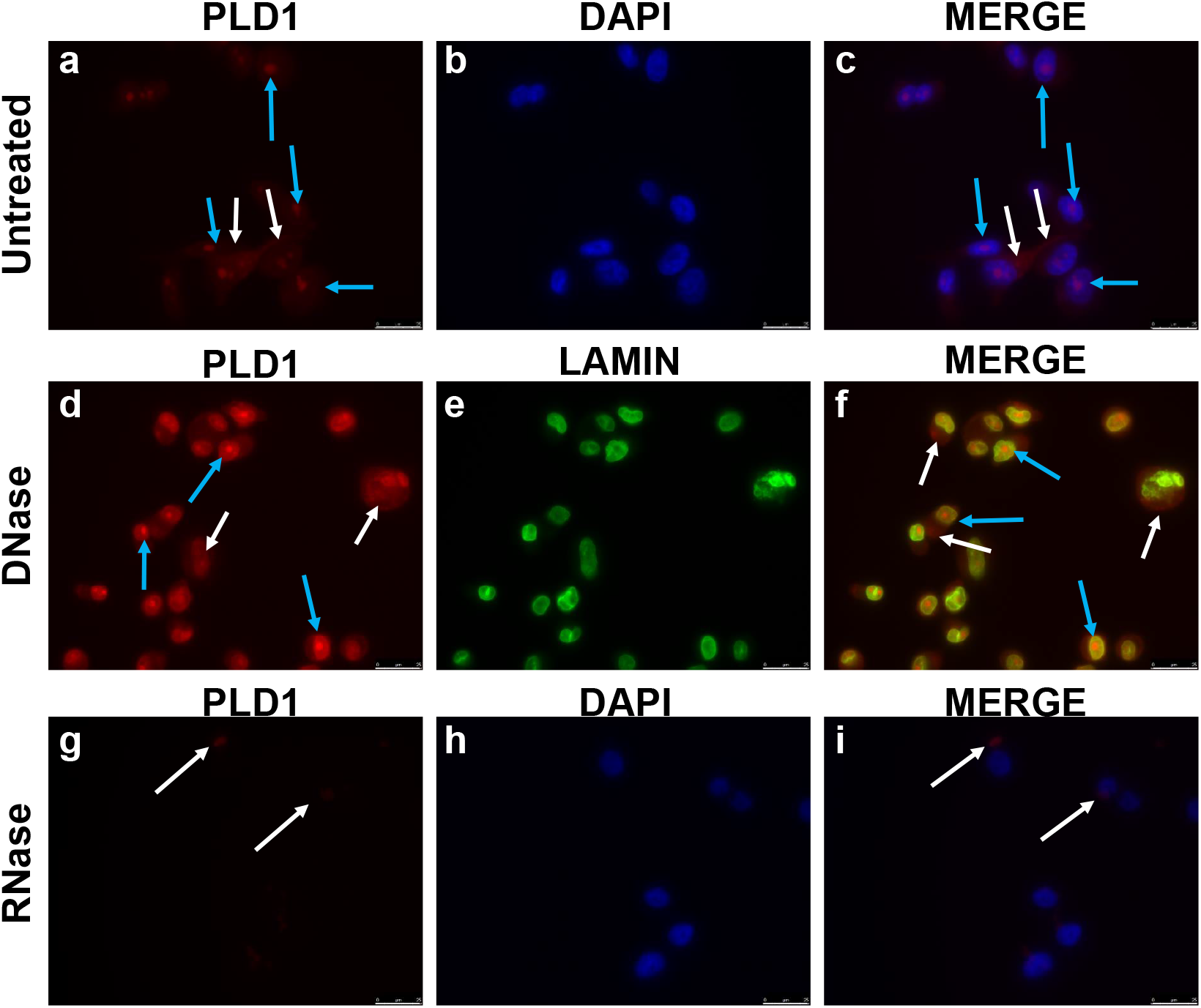
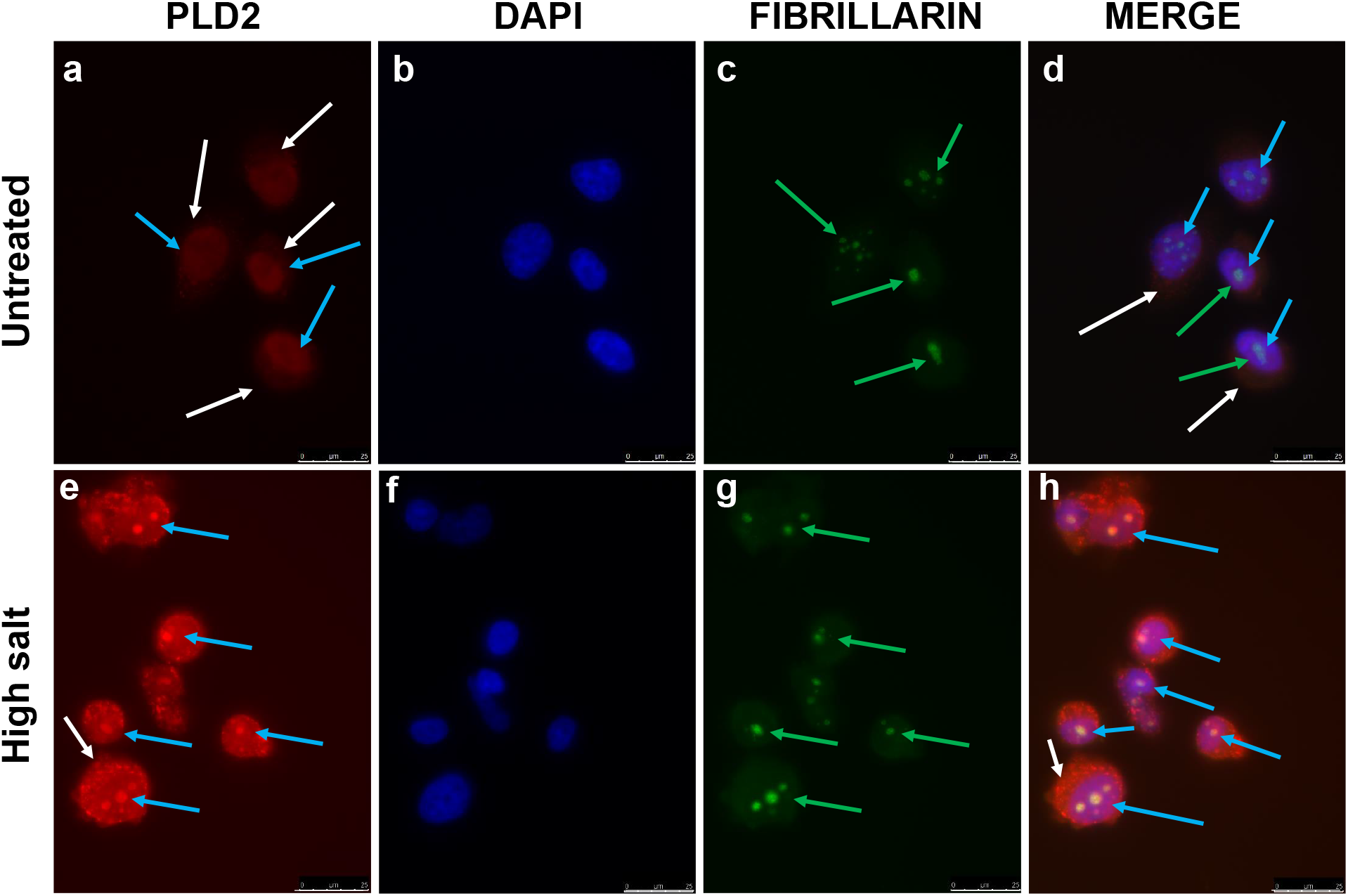
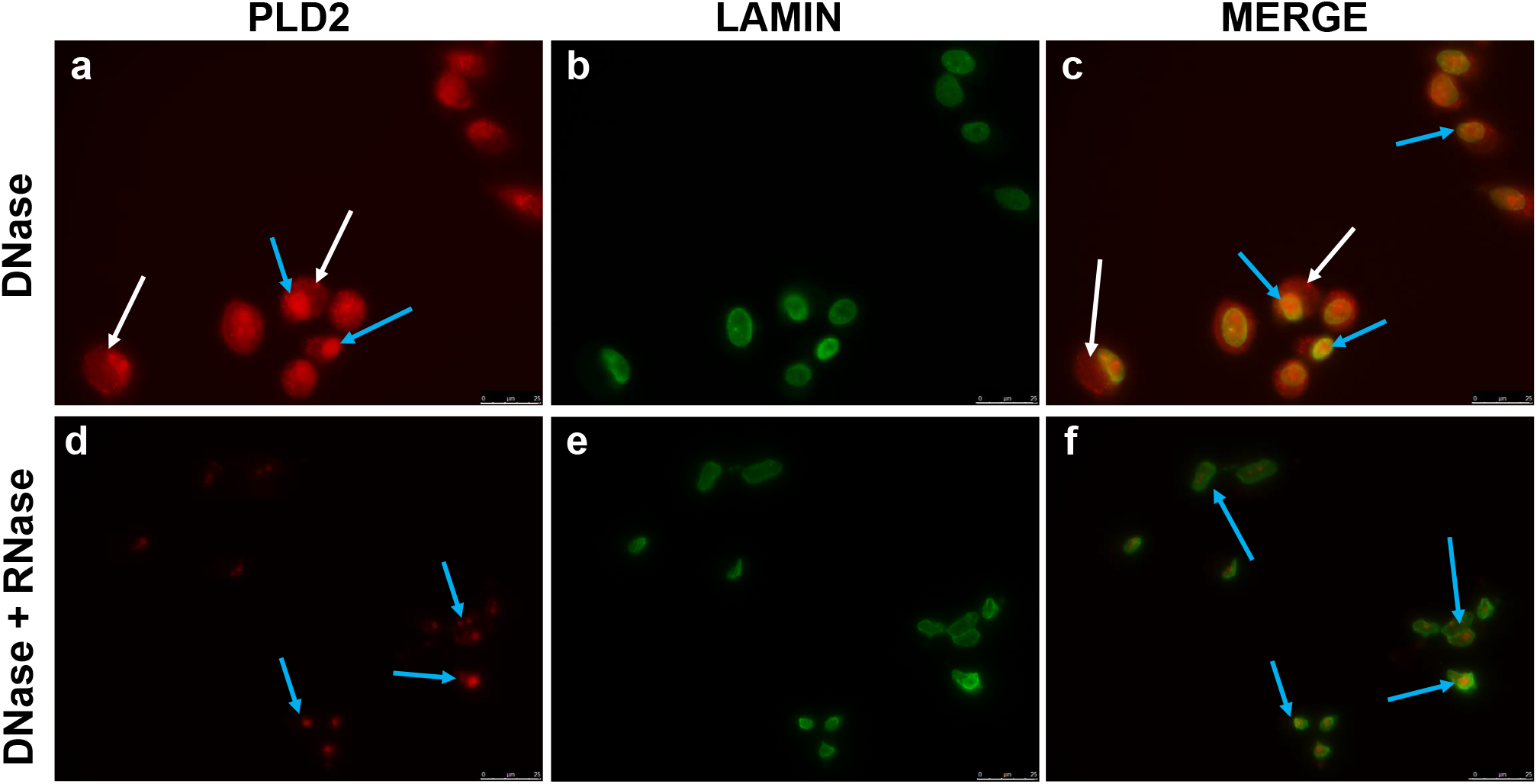
PLD1 and PLD2 in PC3 cells. (**A**), PLD1 protein is detected in the nuclei of PC3 prostate cancer cells after detergent, high salt and DNase treatment but is lost after RNase digestion, red = PLD1, blue = DAPI to detect chromatin, green = lamin to define the perimeter of the nucleus. (**B**), Nuclear PLD2 in PC3 cells is revealed after’ detergent treatment and high salt extraction, and colocalises with fibrillarin, a nucleolus marker, red = PLD2, blue = DAPI, green = fibrillarin. (**C**), some nuclear PLD2 resists subsequent DNase + RNase treatment, red = PLD2, green = Lamin. Scale bar 25 μm. See Methodology for details.

Subsequent digestion of detergent + high salt-treated cells with DNase (Fig. 2C) did not change the detection of PLD2 protein in perinuclear (a, white arrows) and nuclear (a, blue arrows) compartments, indicating that it is not retained by association with chromatin. Inside the nuclei, PLD2 was again detected as a concentrated spot (blue arrows). The overlay with lamin to define the NL (c) confirms PLD2 both in a diffuse perinuclear (white arrows) location and concentrated within nuclei (blue arrows). Further treatment with RNase resulted in the loss of all perinuclear PLD2 protein. However, this RNase treatment did not release the concentrated spots of nuclear PLD2 which appear collapsed in response to digestion of nucleolar RNA but, in the overlay with lamin, remain concentrated within the nucleus (f, blue arrows).

### 3.2. PLD1 and PLD2 in patient-derived PCa cells

In untreated patient-derived PCa cells, PLD1 (Fig. 3A, a) was weakly detected in the cytosol (white arrows) and occasional nuclei (blue arrows) as is apparent in a merged PLD1/DAPI image (3A,c). Initial detergent treatment exposed PLD1 in the nucleus as a concentrated spot (3A,d, blue arrows), clearly shown in the PLD1/DAPI overlay (3A,f). Nuclear PLD1 survives the high salt extraction as shown when the images are merged (3A,i). After DNase treatment PLD1 appeared diffusely distributed within nuclei, at the nuclear membrane (Fig. 3B, a, white arrows) and as concentrated spots (3B, a, blue arrows) within the NL as defined by lamin staining in the merged images (3B,c, blue arrows). Some PLD1 protein was still weakly detected at the nuclear membrane (3B,c, white arrows). Final treatment with RNase removed all PLD1 protein from what remained of the extracted cells (3B, d).

**Figure 3.**
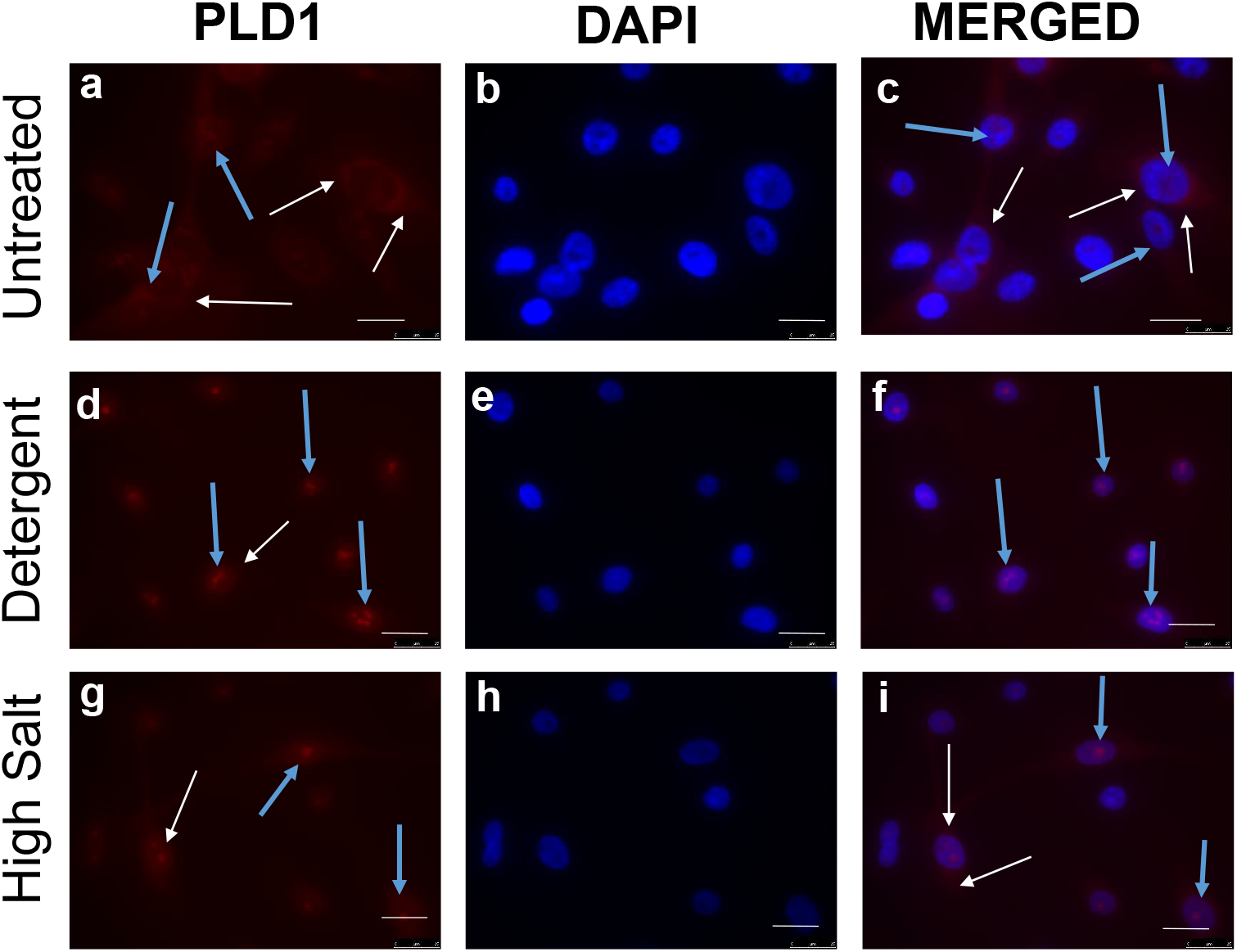
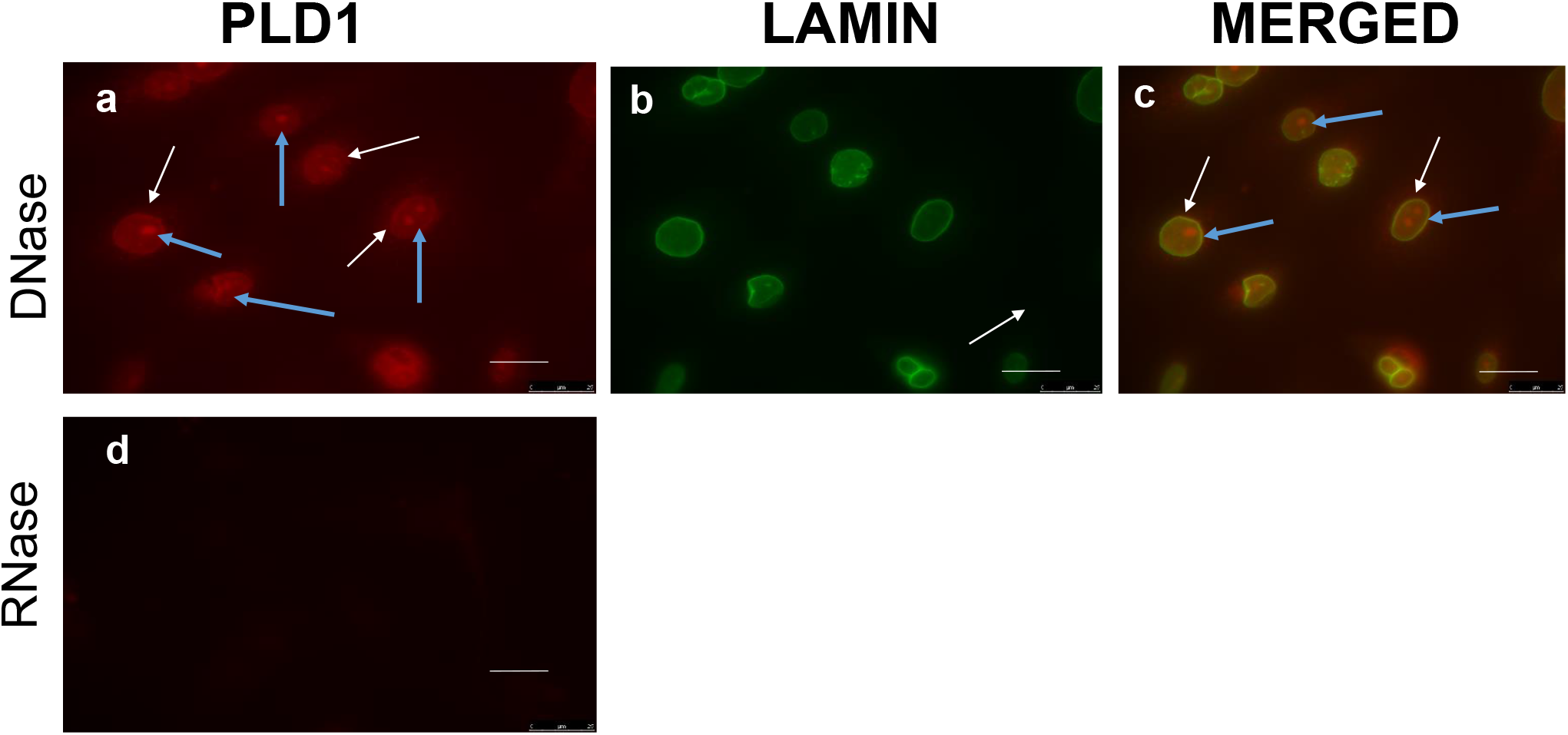
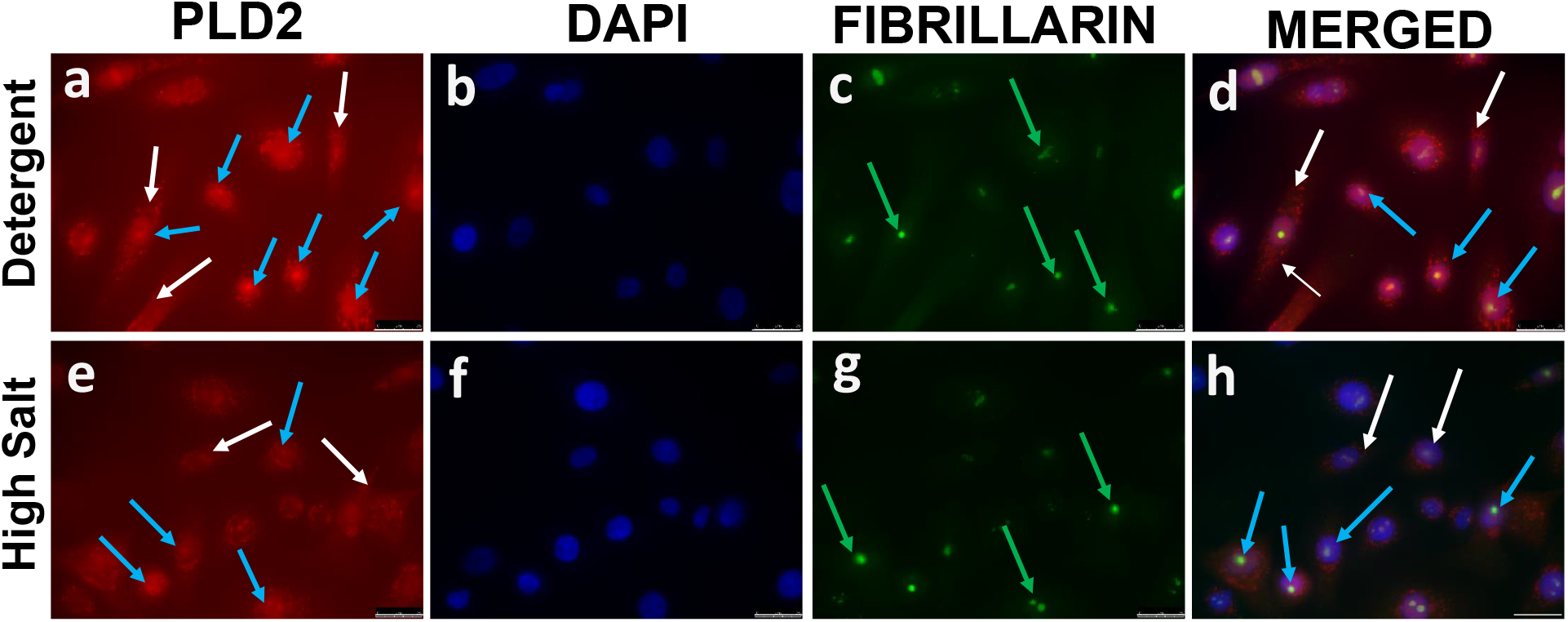
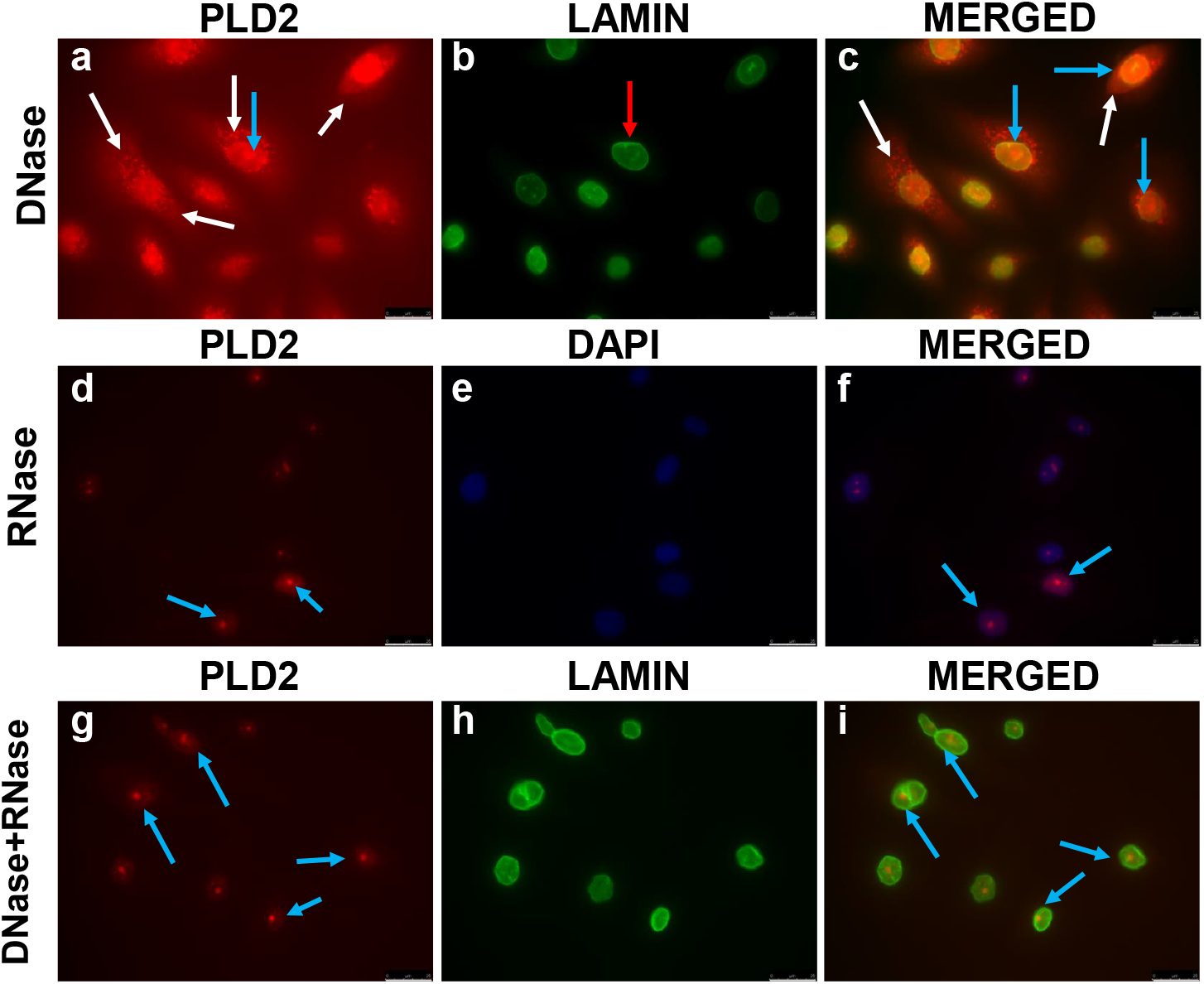
PLD1 and PLD2 in patient-derived prostate cancer cells (H801). (**A**) + (**B**), PLD1 survives extraction as speckles inside the nucleus but is lost after RNase treatment. Red =PLD1, green = Lamin, blue = DAPI. (**C**), PLD2 (red) co-localises with fibrillarin (green). (**D**), PLD2 (red) survives extraction until RNase treatment when diffuse staining in the cytosol is lost but some PLD2 persists in the nucleus. Lamin = green. Scale bar 25 μm. See Methodology for practical detail.

Weak PLD2 expression was detected in untreated patient-derived cells in both cytosol and nuclei (result not shown), similar to PLD2 in untreated PC3 cells (Fig. 2B, a). As before, detergent treatment (Fig. 3C,a) exposed PLD2 as punctate speckles both in a perinuclear location in the cytosol (white arrows) and concentrated in the nucleus (blue arrows). When PLD2:DAPI and fibrillarin images are merged (Fig 3C,d), PLD2 is shown to be colocalised with the (green) fibrillarin nucleolar marker as yellow/white spots (blue arrows). Some PLD2 remained in the cytosol (3C, d, white arrows). This distribution was unchanged on subsequent high salt treatment where again PLD2 co-localised with fibrillarin as revealed by the bright spots in the merged images (Fig 3C, h, blue arrows). DNase treatment (Fig. 3D, a) exposed PLD2 both in the cytosol (white arrows) with a punctate perinuclear distribution and concentrated (blue arrows) within nuclei, as defined by lamin staining (3D, b). This distribution is observed in the merged PLD2/lamin image (3D,c where some PLD2 appears localised at the nuclear membrane (blue arrows). All cytoplasmic and some diffuse nuclear PLD2 staining was lost after RNase treatment or on DNase + RNase treatment together (3D,d, g) but a discrete spot of PLD2 was still detected in what remained of nuclei (3D, d,f,g,i) as identified by lamin staining. Discrete spots of PLD2 staining are clearly observed within the remnants of the nuclei. This result for PLD2 is different from PLD1 staining where RNase treatment removed all PLD1 protein.

### 3.3 Raptor and Rictor expression in the nuclei of patient-derived prostate cancer cells

IF results (Fig. 4.) reveal that mTOR is more strongly expressed in the cytosol of patient-derived PCa cells compared with the nucleus while the opposite is true for expression of Raptor and especially for Rictor. In Fig. 4i blue arrows highlight Rictor concentrated at the plasma membrane of some PCa cells.

**Figure 4.**
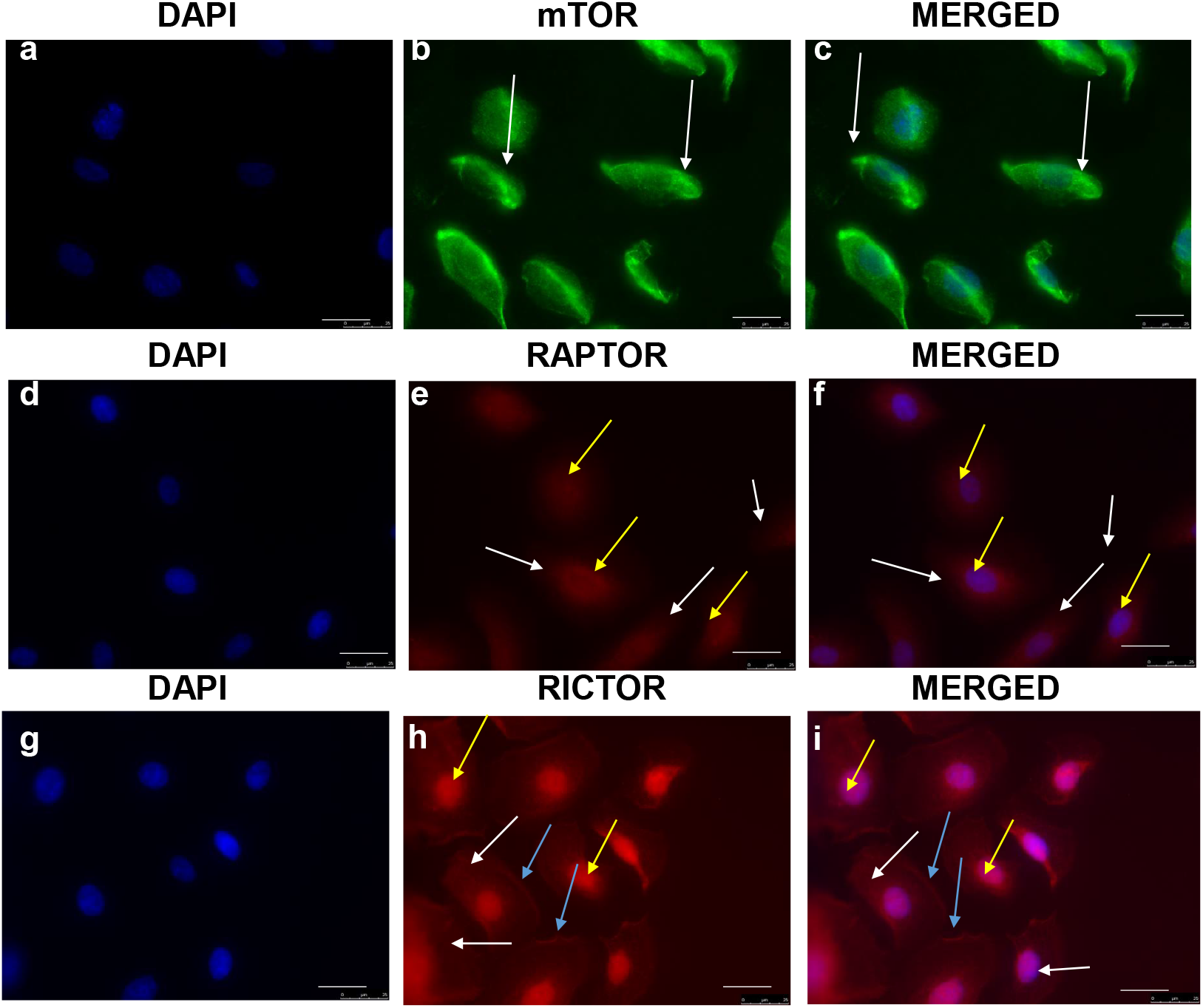
mTOR, Raptor and Rictor protein expression in patient-derived prostate cancer cells. White, yellow and blue arrows define localisation in the cytosol, nuclei and at the plasma membrane respectively (where appropriate). Patient-derived PCa cells were isolated from biopsy tissue (H816) and cultured as described in the Methods section. Scale bar 25 μm.

### 3.4. Effects of inhibiting PLD1 and PLD2 on cell cycle progression

EdU+ cells cycling into or through S phase, cells in G1/G2 and subG1 dying cells with fragmented DNA were identified by the gating strategy from the EdU Lin Area / PI log scatterplot as shown in Fig. 5A for a typical experiment with PC3 cells after 48 hours and Figs. 5B and 5C for two different samples of patient-derived cells (H741, H742) analysed after 72 hours. The important EdU+ cells entering or cycling through S phase were clearly defined. A population of G2/M phase cells was readily identified from non-cycling G1 phase cells with the PC3 cell line (Fig. 5A) but was not detected with the heterogeneous patient-derived cells (Fig. 5B, C). For standardization G2/M cells were combined with G1 phase cells in a non-cycling G1+G2/M group even though we appreciate that these cells are not in the same phase of the cell cycle. Quantification of the cell distribution is shown as percentage values within each gate.

**Figure 5.**
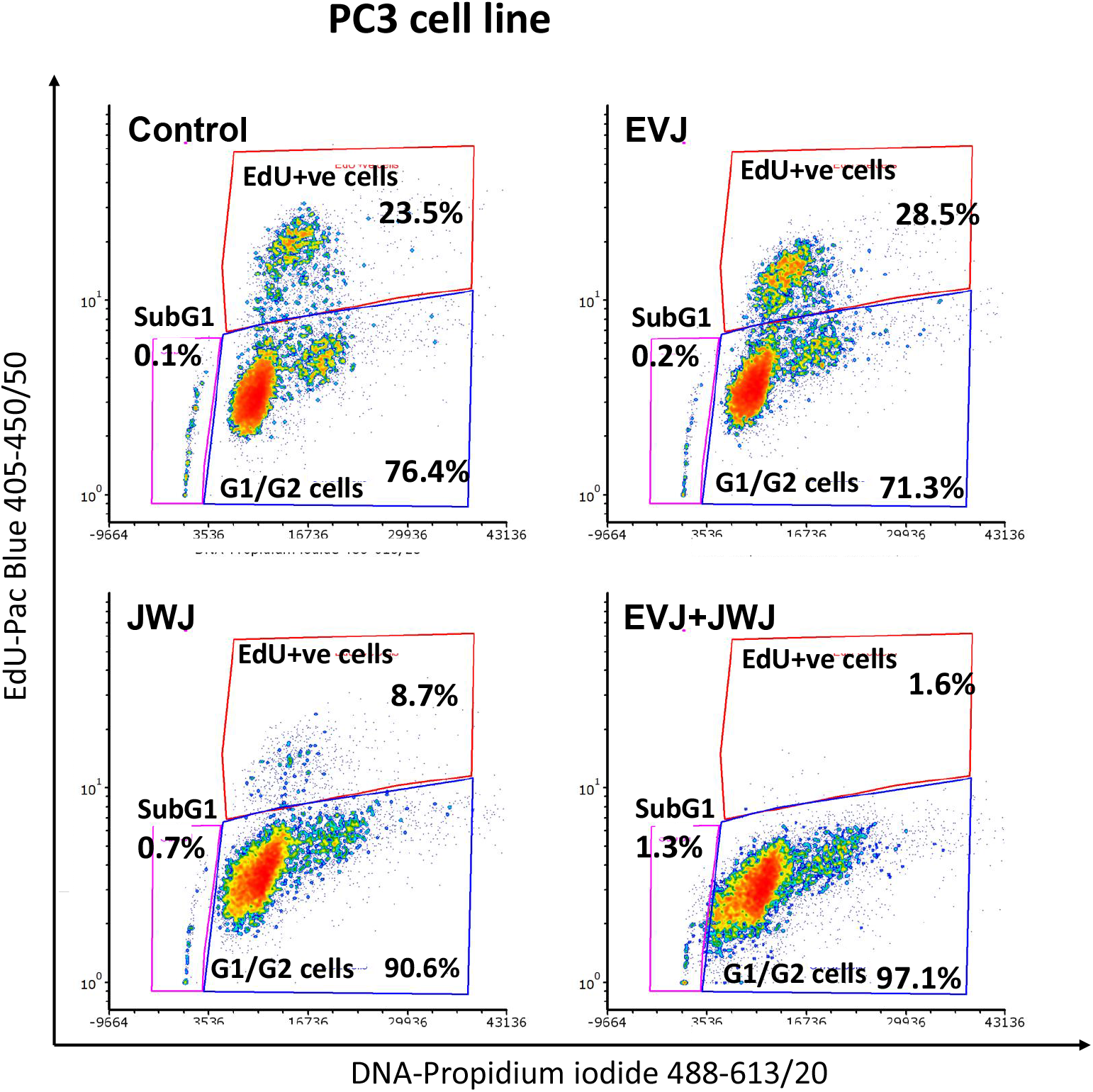
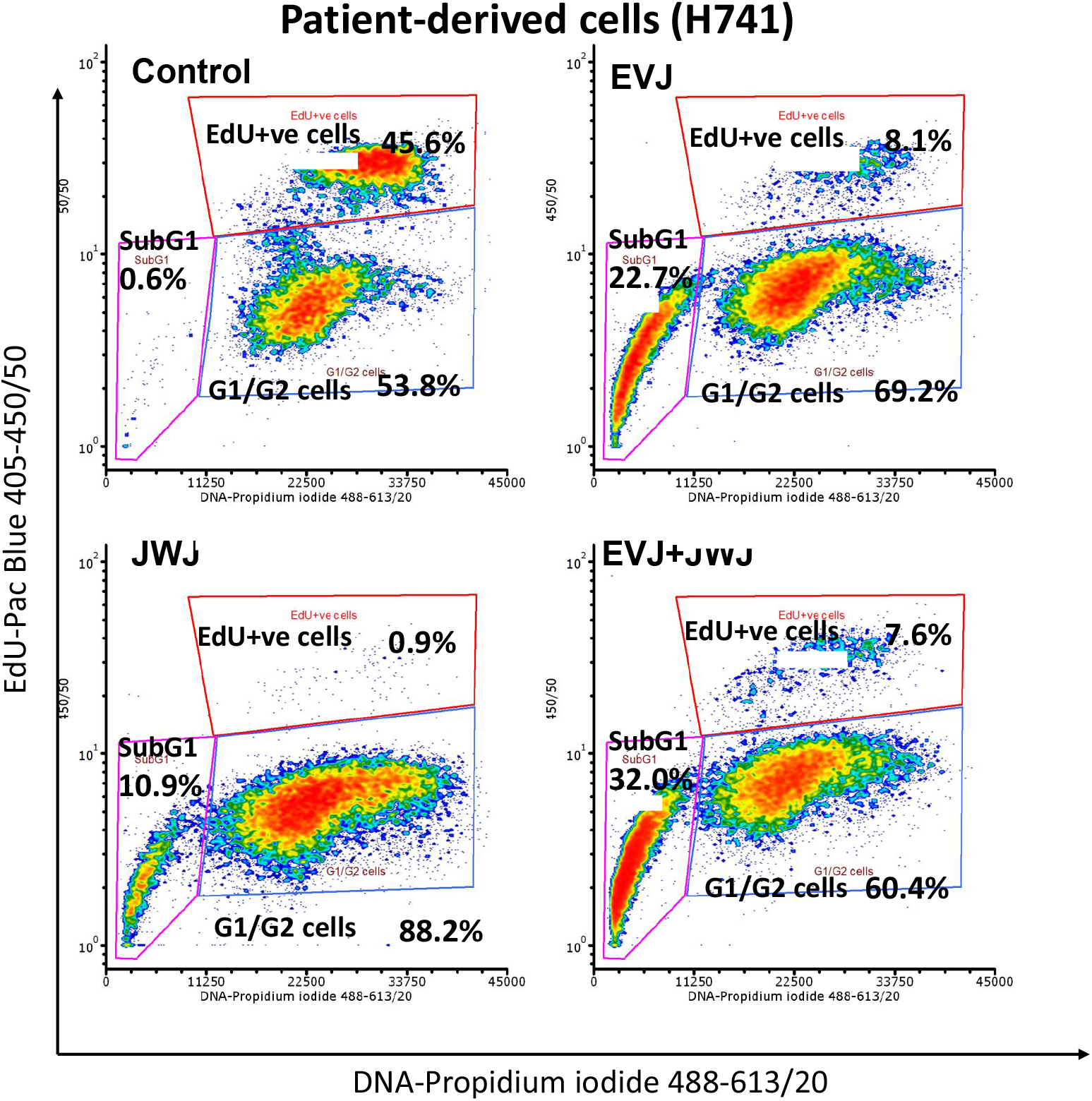
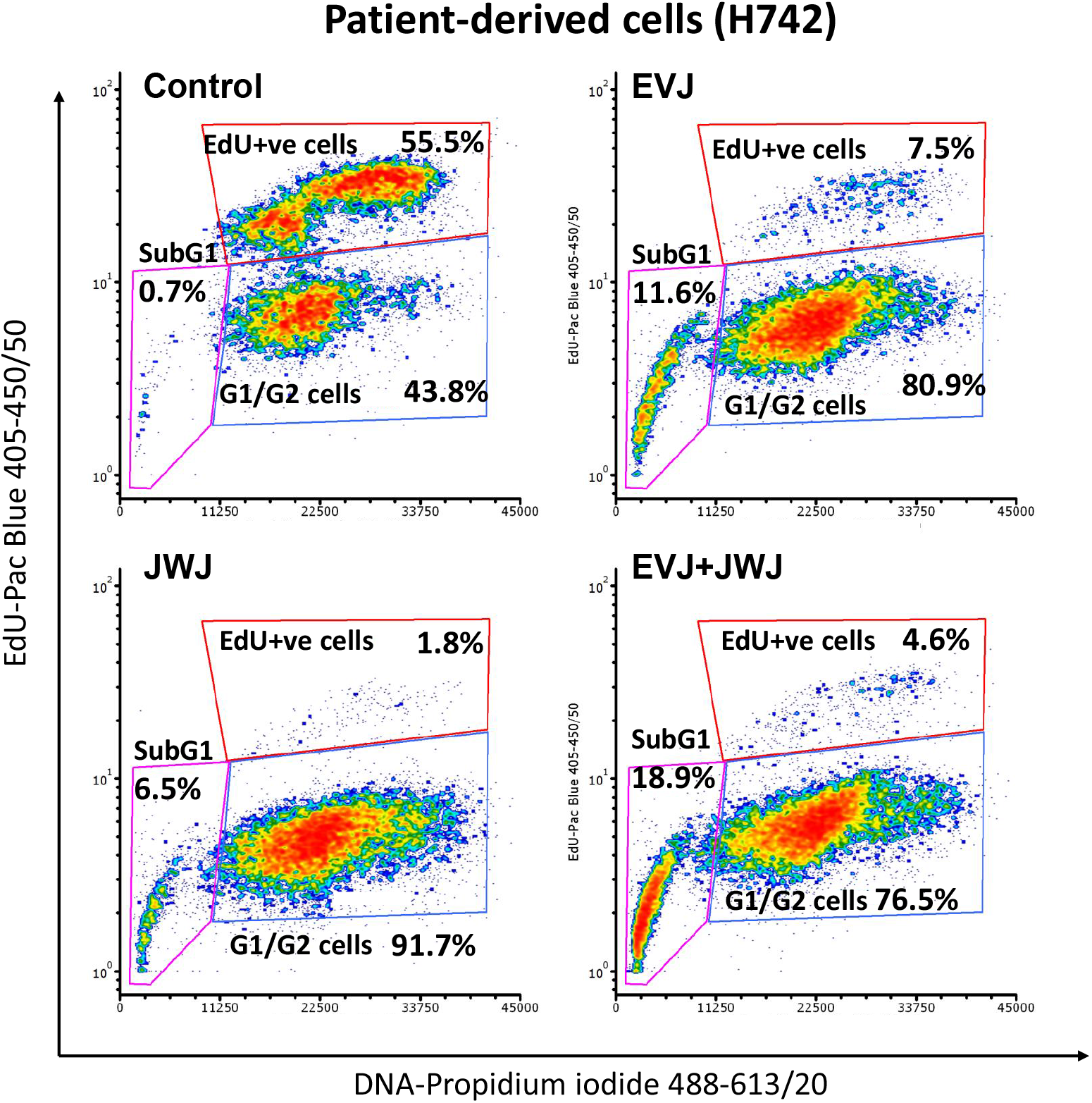
Inhibition of PLD1 or PLD2 decreases PCa cell cycling. (**A**), PC3 cells. (**B**), Patient-derived PCa cells (H741) and (**C**), Patient-derived PCa cells (H742). A third sample of patient-derived cells gave similar results (not shown). The percentage figures in each gate gives the proportion of cells cycling into or passing through S phase, cells in G1+G2/M phases of the cell cycle and, where identified, dying cells with fragmented DNA in a subG1 population. See Methodology for details.

Treatment with the PLD1 inhibitor EVJ at 10 M had virtually no effect on the percentage of PC3 cells cycling in the last 4 hours of the treatment period relative to untreated control cells (Fig. 5 A). However, the PLD2 inhibitor JWJ at 10 M markedly decreased the percentage of PC3 cells entering or passing through S phase from a control value of 23.5% to 8.7% (Fig. 5A). When applied in combination for 48 hours, EVJ + JWJ reduced the percentage of cycling EdU+ PC3 cells even further from a control value of 23.5% to 1.6%; a small (1.3%) subG1 population of dying PC3 cells was also observed.

The effects of EVJ and JWJ on the cell cycle of two separate samples of patient-derived PCa cells (H741, H742) were examined after 72 hours (Figs. 5B, C) since primary cells cycle more slowly than cell lines. Unlike its effects on PC3 prostate cells, treatment of patient-derived cells with the PLD1 inhibitor EVJ caused a marked decrease in the percent of EdU+ cycling cells from control values of 45.6% in H741 and 55.5% in H742 down to 8.1% and 7.5% respectively. Conversely, the percentage of cells between 48 and 72 hours that remained in the combined G1/G2 group and did not enter or pass through S phase was greater in EVJ-treated cells (69.2% in H741, 80.9% in H742) compared with untreated cells (53.8% in H741, 43.8% in H742). Most notably, a subpopulation (8.1%/7.5%, H741/H742) of EdU+ patient-derived PCa cells were resistant to PLD1 inhibition and still entered or passed through S phase. The PLD2 inhibitor JWJ produced a dramatic effect on PCa cell cycling with as few as 0.9% or 1.8% EdU+ cells being detected during the same period. Treatment of either primary cell samples with EVJ and more especially with JWJ increased the percentage of cells in the subG1 population of dying cells with fragmenting DNA. With EVJ and JWJ in combination at half their IC_50_ concentrations a subpopulation of EdU+ patient-derived PCa cells still entering into or passing through S phase (7.6% in H741, 4.6% in H742) was detected, rather similar to the population of EdU+ cells detected with EVJ alone.

## 4. Discussion

The unique and scarce patient-derived primary PCa cells used here, as in our earlier work^9,24^, have a basal cell phenotype (CK5^+^ / CK8^-^ / AR^-^ / CD49b^+^ / CD44^+^ / CD24^+^)^45,46^. These are the main cells cultured from PCa biopsy tissue because tumorigenic luminal cells which account for the bulk of PCa tissue do not adapt to culture as they are terminally differentiated. Our basal PCa cells can be differentiated to a luminal phenotype in culture^47^ and a significant proportion have the TMPRSS2:ERG fusion gene (if present in the original cancer), a feature of up to 50% of prostate cancers^48^. They are more invasive than BPH cell cultures^49^, have increased expression of the cancer cell-associated protein POLR3G^50^ and are more resistant to chemotherapy and radio-therapy treatment than other prostate cells^51^. The small yield of cells obtained per biopsy means that experiments cannot usually be replicated. We used PC3 cells for comparison since they do not express AR, like the patient-derived PCa basal cells. 22RV1 cells would have been an alternative choice but they express AR^52^.

The extraction protocol exposed epitopes on nuclear PLD1 and nuclear PLD2 that are normally masked as noted for other proteins^35-37^. This is shown by the enhanced fluorescence detected after treating cells with CSK buffer-detergent alone (Fig. 3A for PLD1), after detergent and high salt (Fig. 2B for PLD2 in PC3 cells) or after detergent, high salt and DNase to remove chromatin (Figs. 2A, 3B, for PLD1). These extraction results confirm and extend our earlier IF and subcellular fractionation/western blot experiments showing that PLD1 and PLD2^9,24^ are partially expressed in prostate cell nuclei, in agreement with other reports^4,25-27,53^. In support, a nuclear phospholipase, PI-PLCβ1 has effects on G2/M progression^54,55^ while the nuclear localisation of a phospholipase A2-α depends on the state of cell proliferation^56^. Like mTOR in the cytosol, chromatin-associated nmTOR^32^ will require PtdOH for activity, providing a rationale for nPLD1 and nPLD2 expression. Much of the PtdCho required by nPLD1 and nPLD2 as a substrate occurs in microdomains with sphingomyelin and cholesterol in the inner nuclear membrane (INM) where cell proliferation is regulated^57-59^. mTOR requires unsaturated species of PtdOH for activation^60^ so at the INM, nPLD1 and/or nPLD2 need to select the appropriate species of PtdCho for hydrolysis. Interestingly, PLD1 and PLD2 seem to differ in the PtdOH species they produce from PtdCho^61^. Endonuclear PtdCho associated with both chromatin and the NM, as well as in complexes as proteolipids^57-59^, is another potential substrate for both nPLD1 and nPLD2. How nPLD1 and nPLD2 are regulated is unknown but in vascular smooth muscle cells nPLD1 is activated by cell surface G-protein-coupled receptors^62^. Nuclear PI-PLCβ1 is also activated by cell surface events^55^. The IF results (Fig.4) clearly reveal that both Raptor and more prominently Rictor, are expressed in PCa cell nuclei so are available to complex with nmTOR as found in the control of mitochondrial gene expression^33^.

Our results indicate that PLD1 and PLD2 are associated with different binding-partners in PCa cell nuclei since PLD1 is released by RNase treatment while nPLD2 is resistant. This is clearly shown by the bright punctate PLD2 fluorescence remaining within the nucleus after DNase and RNase treatments (eg. Figs. 2C, 3D). At such different binding sites/locations nPLD1 and nPLD2 may have access to different species of PtdCho so that the nature of the PtdOH species generated subsequently determines their differential functions in the nucleus^61,63^. The identification of these nuclear PLD binding sites may be complicated by the fact that the protein composition of the NM can change during disease progression^64,65^.

It is notable in Figs. 2B and 3B that the distinct spots of PLD2 fluorescence match fluorescence from the nucleolar marker fibrillarin^66^. Intriguingly, Foster and colleagues have reported a link between PLD1 or PLD2 and the nucleolar stress pathway^67^ whereby increased expression of PLD1 or PLD2 raises levels of the nucleolar oncoprotein MDM2 resulting in an increased turnover of the tumour suppressor protein p53. This effect involves mTOR and Raf, which need PtdOH for activity^16,17,68^ and membrane binding^69^ respectively. Such a study provides a link between PLD2 and the nucleolus^70^, since in non-tumorigenic cells when MDM2 is associated with ARF in the nucleolus, p53 activity is maintained at a low level^71^. Later work has revealed that PLD stabilization of HDM2 occurs through mTORC2^72^ which contains Rictor as expressed in PCa cell nuclei (Fig 4.).

Our results with EdU labelling to define replicating cells reveal that both PLD1 and PLD2 contribute to the control of cycling by patient-derived PCa cells. This is in agreement with findings on colorectal cells and fibroblasts^10,30^ and also that overexpression of both PLD1 and PLD2 overcomes cell cycle arrest induced by high intensity Raf signalling^28^. The data confirm that PLD2, and to a lesser extent PLD1, contribute significantly to the function of mTOR in its control of the cell cycle^17^ in patient-derived primary PCa cells, and explain why inhibiting PLD1 and especially PLD2 blocks PCa cell proliferation so effectively^9,24^. PLD2 can also influence the entry or passage through the cell cycle by PCa cells, mediated by cyclin D3, an activator of the G1 to S phase transition^10^, and (with PLD1) by inhibiting expression of the CDK inhibitor p21 gene^29^. The finding that a greater proportion of patient-derived cells are in a subG1 population after treatment with EVJ or JWJ compared with PC3 cells is not surprising since the patient-derived cells were in contact with the inhibitors for longer. EVJ and JWJ were used in the 6-11 μM range after Lavieri et al^43,44^ and Mathews et al^73^ because >95% of these lipophilic inhibitors bind to serum- or supplement-derived proteins in the growth medium and because they then have to cross the plasma membrane of the cultured cells. The selective effects of these inhibitors on PLD isoforms in intact cells are maintained at these concentrations as reported by others^44,73-75^ and as shown by the fact that EVJ and JWJ produce markedly different effects on cell cycling at the concentrations used.

Our finding that inhibiting PLD1 induced a substantial decrease in the percentage of cycling EdU+ patient-derived PCa cells yet had no effect on PC3 cells was unexpected and may reflect an interesting difference on the role of PLD1 in primary patient-derived cells and prostate cell lines. Unlike the relatively homogeneous cellular content of established prostate cell lines, each preparation of patient-derived cells is a heterogeneous population unique to each patient^76^. Additionally, differences in PLD1 expression^24^ and/or unknown factors in foetal calf serum and in the supplement added to the primary PCa cell medium might be responsible. Our finding (Fig. 5B, C) that a small subpopulation of patient-derived cells continue cycling when EVJ and JWJ are used in combination is probably due to the incomplete inhibition of PLD2 as when in combination EVJ and JWJ were used at half their IC_50_ values to minimize cell death with the longer incubation period used. The results still reveal that PLD2 is the important isoform to target in any therapeutic use of these two PLD inhibitors. The antipsychotic agent halopemide on which the PLD inhibitors are based, has been tested in clinical trials and is well tolerated strengthening the case for making the PLD2 inhibitor a potential therapeutic agent^77,78^.

**Supplementary Figure:** Figure S1: Identification of PLD1 in PC3 cells at each stage of the nuclear matrix extraction procedure.

## Author Contributions

MGR took the lead in writing the manuscript. ARN carried out most of the experiments and with FF prepared the ICC figures. KH analysed the cell cycle data and prepared the associated figures. SB provided the PLD2 antibody and advised on its use. DC provided expertise and advice on the nuclear matrix extraction procedure. LA carried out some of the cell cycle experiments. NJM guided the research throughout the project. All authors read the manuscript and provided feedback.

## Informed Consent

All prostate tissue samples were obtained with patient consent and full ethical approval (South Yorkshire Research Ethics Committee, Yorkshire and the Humber, REC:07/H1304/121). There is no identifying patient information in this publication. This study was performed in accordance with the Declaration of Helsinki.

## Consent for publication

All authors confirm that this article can be considered for publication.

## Data availability

The data used in this article is stored securely at the University of York and is available on request.

## Acknowledgements

Hannah Walker provided excellent technical support. We thank the Coverley group for assistance and surgeons and patients from Castle Hill Hospital, Cottingham, Hull (principally Mr Matthew Simms) and Dr Vincent Mann for providing prostate biopsy tissue.

## References

1. Uchida N, Okamura S, Kuwano H. Phospholipase D activity in human gastric carcinoma. Anticancer research 1999; 19(1B): 671–675.

2. Uchida N, Okamura S, Nagamachi Y, Yamashita S. Increased phospholipase D activity in human breast cancer. Journal of cancer research and clinical oncology 1997; 123(5): 280–285.

3. Noh D-Y, Ahn S-J, Lee R-A, Park I-A, Kim J-H, Suh P-G et al. Overexpression of phospholipase D1 in human breast cancer tissues. Cancer Letters 2000; 161(2): 207-214; doi http://dx.doi.org/10.1016/S0304-3835(00)00612-1.

4. Zhao Y, Ehara H, Akao Y, Shamoto M, Nakagawa Y, Banno Y et al. Increased Activity and Intranuclear Expression of Phospholipase D2 in Human Renal Cancer. Biochemical and Biophysical Research Communications 2000; 278(1): 140-143; doi http://dx.doi.org/10.1006/bbrc.2000.3719.

5. Saito M, Iwadate M, Higashimoto M, Ono K, Takebayashi Y, Takenoshita S. Expression of phospholipase D2 in human colorectal carcinoma. Oncology reports 2007; 18(5): 1329–1334.

6. Kim Y-R, Byun HS, Won M, Park KA, Kim JM, Choi BL et al. Modulatory role of phospholipase D in the activation of signal transducer and activator of transcription (STAT)-3 by thyroid oncogenic kinase RET/PTC. BMC Cancer (journal article) 2008; 8(1): 144; doi 10.1186/1471-2407-8-144.

7. Henkels KM, Boivin GP, Dudley ES, Berberich SJ, Gomez-Cambronero J. Phospholipase D (PLD) drives cell invasion, tumor growth and metastasis in a human breast cancer xenograph model. Oncogene 2013; 32(49): 5551–5562; doi 10.1038/onc.2013.207.

8. Kandori S, Kojima T, Matsuoka T, Yoshino T, Sugiyama A, Nakamura E et al. Phospholipase D2 promotes disease progression of renal cell carcinoma through the induction of angiogenin. Cancer Sci 2018; 109(6): 1865-1875; e-pub ahead of print 2018/04/17; doi 10.1111/cas.13609.

9. Noble AR, Hogg K, Suman R, Berney DM, Bourgoin S, Maitland NJ et al. Phospholipase D2 in prostate cancer: protein expression changes with Gleason score. British journal of cancer 2019; 121(12): 1016-1026; e-pub ahead of print 2019/11/02; doi 10.1038/s41416-019-0610-7.

10. Min DS, Kwon TK, Park WS, Chang JS, Park SK, Ahn BH et al. Neoplastic transformation and tumorigenesis associated with overexpression of phospholipase D isozymes in cultured murine fibroblasts. Carcinogenesis 2001; 22(10); doi 10.1093/carcin/22.10.1641.

11. Zheng Y, Rodrik V, Toschi A, Shi M, Hui L, Shen Y et al. Phospholipase D couples survival and migration signals in stress response of human cancer cells. The Journal of biological chemistry 2006; 281(23): 15862–15868; doi 10.1074/jbc.M600660200.

12. Utter M, Chakraborty S, Goren L, Feuser L, Zhu YS, Foster DA. Elevated phospholipase D activity in androgen-insensitive prostate cancer cells promotes both survival and metastatic phenotypes. Cancer Lett 2018; 423: 28–35; e-pub ahead of print 2018/03/11; doi 10.1016/j.canlet.2018.03.006.

13. Borel M, Cuvillier O, Magne D, Mebarek S, Brizuela L. Increased phospholipase D activity contributes to tumorigenesis in prostate cancer cell models. Molecular and Cellular Biochemistry 2020; 473(1): 263–279; doi 10.1007/s11010-020-03827-2.

14. Borel M, Lollo G, Magne D, Buchet R, Brizuela L, Mebarek S. Prostate cancer-derived exosomes promote osteoblast differentiation and activity through phospholipase D2. Biochimica et Biophysica Acta (BBA) - Molecular Basis of Disease 2020; 1866(12): 165919; doi https://doi.org/10.1016/j.bbadis.2020.165919.

15. Fang Y, Vilella-Bach M, Bachmann R, Flanigan A, Chen J. Phosphatidic acid-mediated mitogenic activation of mTOR signaling. Science (New York, NY) 2001; 294(5548): 1942-1945; e-pub ahead of print 2001/12/01; doi 10.1126/science.1066015.

16. Foster DA, Salloum D, Menon D, Frias MA. Phospholipase D and the maintenance of phosphatidic acid levels for regulation of mammalian target of rapamycin (mTOR). The Journal of biological chemistry 2014; 289(33): 22583–22588; doi 10.1074/jbc.R114.566091.

17. Foster DA. Phosphatidic acid signaling to mTOR: signals for the survival of human cancer cells. Biochimica et biophysica acta 2009; 1791(9): 949–955; doi 10.1016/j.bbalip.2009.02.009.

18. Toschi A, Lee E, Xu L, Garcia A, Gadir N, Foster DA. Regulation of mTORC1 and mTORC2 complex assembly by phosphatidic acid: competition with rapamycin. Molecular and cellular biology 2009; 29(6): 1411–1420; e-pub ahead of print 2008/12/31; doi 10.1128/mcb.00782-08.

19. Foster DA. Phosphatidic acid and lipid-sensing by mTOR. Trends in endocrinology and metabolism: TEM 2013; 24(6): 272–278; doi 10.1016/j.tem.2013.02.003.

20. Kim Y, Han JM, Han BR, Lee KA, Kim JH, Lee BD et al. Phospholipase D1 is phosphorylated and activated by protein kinase C in caveolin-enriched microdomains within the plasma membrane. The Journal of biological chemistry 2000; 275(18): 13621–13627; e-pub ahead of print 2000/05/02.

21. Hu T, Exton JH. Mechanisms of regulation of phospholipase D1 by protein kinase Calpha. The Journal of biological chemistry 2003; 278(4): 2348–2355; e-pub ahead of print 2002/11/15; doi 10.1074/jbc.M210093200.

22. Du G, Huang P, Liang BT, Frohman MA. Phospholipase D2 localizes to the plasma membrane and regulates angiotensin II receptor endocytosis. Molecular biology of the cell 2004; 15(3): 1024–1030; e-pub ahead of print 2004/01/14; doi 10.1091/mbc.e03-09-0673.

23. Bruntz RC, Taylor HE, Lindsley CW, Brown HA. Phospholipase D2 mediates survival signaling through direct regulation of Akt in glioblastoma cells. The Journal of biological chemistry 2014; 289(2): 600–616; e-pub ahead of print 2013/11/22; doi 10.1074/jbc.M113.532978.

24. Noble AR, Maitland NJ, Berney DM, Rumsby MG. Phospholipase D inhibitors reduce human prostate cancer cell proliferation and colony formation. British journal of cancer 2018; 118(2): 189–199; e-pub ahead of print 2017/11/15; doi 10.1038/bjc.2017.391.

25. Freyberg Z, Sweeney D, Siddhanta A, Bourgoin S, Frohman M, Shields D. Intracellular localization of phospholipase D1 in mammalian cells. Molecular biology of the cell 2001; 12(4): 943–955; e-pub ahead of print 2001/04/11.

26. Freyberg Z, Bourgoin S, Shields D. Phospholipase D2 is localized to the rims of the Golgi apparatus in mammalian cells. Molecular biology of the cell 2002; 13(11): 3930–3942; e-pub ahead of print 2002/11/14; doi 10.1091/mbc.02-04-0059.

27. Jang YH, Min DS. The hydrophobic amino acids involved in the interdomain association of phospholipase D1 regulate the shuttling of phospholipase D-1 from vesicular organelles into the nucleus. Experimental and Molecular Medicine 2012; 44(10): 571–577; doi 10.3858/emm.2012.44.10.065.

28. Joseph T, Bryant A, Frankel P, Wooden R, Kerkhoff E, Rapp UR et al. Phospholipase D overcomes cell cycle arrest induced by high-intensity Raf signaling. Oncogene (Short Report) 2002; 21: 3651; doi 10.1038/sj.onc.1205380.

29. Kwun HJ, Lee JH, Min DS, Jang KL. Transcriptional repression of cyclin-dependent kinase inhibitor p21 gene by phospholipase D1 and D2. FEBS Lett 2003; 544(1-3): 38–44; e-pub ahead of print 2003/06/05; doi 10.1016/s0014-5793(03)00446-0.

30. Baan B, Dihal AA, Hoff E, Bos CL, Voorneveld PW, Koelink PJ et al. 5-Aminosalicylic acid inhibits cell cycle progression in a phospholipase D dependent manner in colorectal cancer. Gut 2012; 61(12): 1708–1715; e-pub ahead of print 2011/12/22; doi 10.1136/gutjnl-2011-301626.

31. Wang X, Proud CG. Nutrient control of TORC1, a cell-cycle regulator. Trends in cell biology 2009; 19(6): 260–267; e-pub ahead of print 2009/05/08; doi 10.1016/j.tcb.2009.03.005.

32. Tsang CK, Liu H, Zheng XF. mTOR binds to the promoters of RNA polymerase I-and III-transcribed genes. Cell cycle (Georgetown, Tex) 2010; 9(5): 953–957; e-pub ahead of print 2009/12/30; doi 10.4161/cc.9.5.10876.

33. Cunningham JT, Rodgers JT, Arlow DH, Vazquez F, Mootha VK, Puigserver P. mTOR controls mitochondrial oxidative function through a YY1–PGC-1α transcriptional complex. Nature 2007; 450(7170): 736–740; doi 10.1038/nature06322.

34. Audet-Walsh E, Dufour CR, Yee T, Zouanat FZ, Yan M, Kalloghlian G et al. Nuclear mTOR acts as a transcriptional integrator of the androgen signaling pathway in prostate cancer. Genes Dev 2017; 31(12): 1228–1242; e-pub ahead of print 2017/07/21; doi 10.1101/gad.299958.117.

35. Stewart ER, Coverley D. Visualization of Hidden Epitopes at the Inactive X Chromosome. Methods in molecular biology (Clifton, NJ) 2018; 1861: 103–112; e-pub ahead of print 2018/09/16; doi 10.1007/978-1-4939-8766-5_9.

36. Wilson RH, Hesketh EL, Coverley D. The Nuclear Matrix: Fractionation Techniques and Analysis. Cold Spring Harb Protoc 2016; 2016(1): pdb.top074518; e-pub ahead of print 2016/01/06; doi 10.1101/pdb.top074518.

37. Wilson RH, Coverley D. Relationship between DNA replication and the nuclear matrix. Genes Cells 2013; 18(1): 17–31; e-pub ahead of print 2012/11/09; doi 10.1111/gtc.12010.

38. Chen J, Shen Y, Jiao R, Zhai Z. Composition and structure of nucleolar skeleton (nucleolar matrix) : Actin and fibrillarin are two main protein components of nucleolar skeleton. Sci China C Life Sci 1999; 42(1): 34–42; e-pub ahead of print 2008/08/30; doi 10.1007/bf02881745.

39. Vlcek S, Dechat T, Foisner R. Nuclear envelope and nuclear matrix: interactions and dynamics. Cellular and Molecular Life Sciences CMLS 2001; 58(12): 1758–1765; doi 10.1007/PL00000815.

40. Scott SA, Selvy PE, Buck JR, Cho HP, Criswell TL, Thomas AL et al. Design of isoform-selective phospholipase D inhibitors that modulate cancer cell invasiveness. Nature chemical biology 2009; 5(2): 108–117; doi 10.1038/nchembio.140.

41. Denmat-Ouisse LA, Phebidias C, Honkavaara P, Robin P, Geny B, Min DS et al. Regulation of constitutive protein transit by phospholipase D in HT29-cl19A cells. The Journal of biological chemistry 2001; 276(52): 48840–48846; e-pub ahead of print 2001/11/01; doi 10.1074/jbc.M104276200.

42. Lewis JA, Scott SA, Lavieri R, Buck JR, Selvy PE, Stoops SL et al. Design and synthesis of isoform-selective phospholipase D (PLD) inhibitors. Part I: Impact of alternative halogenated privileged structures for PLD1 specificity. Bioorganic & medicinal chemistry letters 2009; 19(7): 1916–1920; e-pub ahead of print 2009/03/10; doi 10.1016/j.bmcl.2009.02.057.

43. Lavieri R, Scott SA, Lewis JA, Selvy PE, Armstrong MD, Alex Brown H et al. Design and synthesis of isoform-selective phospholipase D (PLD) inhibitors. Part II. Identification of the 1,3,8-triazaspiro[4,5]decan-4-one privileged structure that engenders PLD2 selectivity. Bioorganic & medicinal chemistry letters 2009; 19(8): 2240–2243; e-pub ahead of print 2009/03/21; doi 10.1016/j.bmcl.2009.02.125.

44. Lavieri RR, Scott SA, Selvy PE, Kim K, Jadhav S, Morrison RD et al. Design, synthesis, and biological evaluation of halogenated N-(2-(4-oxo-1-phenyl-1,3,8-triazaspiro[4.5]decan-8-yl)ethyl)benzamides: discovery of an isoform-selective small molecule phospholipase D2 inhibitor. Journal of medicinal chemistry 2010; 53(18): 6706–6719; e-pub ahead of print 2010/08/26; doi 10.1021/jm100814g.

45. Frame FM, Pellacani D, Collins AT, Maitland NJ. Harvesting Human Prostate Tissue Material and Culturing Primary Prostate Epithelial Cells. Methods in molecular biology (Clifton, NJ) 2016; 1443: 181–201; e-pub ahead of print 2016/06/02; doi 10.1007/978-1-4939-3724-0_12.

46. Pellacani D, Droop AP, Frame FM, Simms MS, Mann VM, Collins AT et al. Phenotype-independent DNA methylation changes in prostate cancer. British journal of cancer 2018; 119(9): 1133–1143; e-pub ahead of print 2018/10/16; doi 10.1038/s41416-018-0236-1.

47. Swift SL, Burns JE, Maitland NJ. Altered expression of neurotensin receptors is associated with the differentiation state of prostate cancer. Cancer research 2010; 70(1): 347–356; e-pub ahead of print 2010/01/06; doi 10.1158/0008-5472.can-09-1252.

48. Polson ES, Lewis JL, Celik H, Mann VM, Stower MJ, Simms MS et al. Monoallelic expression of TMPRSS2/ERG in prostate cancer stem cells. Nat Commun 2013; 4: 1623; e-pub ahead of print 2013/03/29; doi 10.1038/ncomms2627.

49. Collins AT, Berry PA, Hyde C, Stower MJ, Maitland NJ. Prospective identification of tumorigenic prostate cancer stem cells. Cancer research 2005; 65(23): 10946–10951; e-pub ahead of print 2005/12/03; doi 10.1158/0008-5472.Can-05-2018.

50. Petrie JL, Swan C, Ingram RM, Frame FM, Collins AT, Dumay-Odelot H et al. Effects on prostate cancer cells of targeting RNA polymerase III. Nucleic Acids Res 2019; 47(8): 3937–3956; e-pub ahead of print 2019/03/02; doi 10.1093/nar/gkz128.

51. Ulukaya E, Frame FM, Cevatemre B, Pellacani D, Walker H, Mann VM et al. Differential cytotoxic activity of a novel palladium-based compound on prostate cell lines, primary prostate epithelial cells and prostate stem cells. PloS one 2013; 8(5): e64278; e-pub ahead of print 2013/05/16; doi 10.1371/journal.pone.0064278.

52. Sampson N, Neuwirt H, Puhr M, Klocker H, Eder IE. In vitro model systems to study androgen receptor signaling in prostate cancer. Endocrine-related cancer 2013; 20(2): R49–64; e-pub ahead of print 2013/03/01; doi 10.1530/ERC-12-0401.

53. Mahankali M, Henkels KM, Speranza F, Gomez-Cambronero J. A non-mitotic role for Aurora kinase A as a direct activator of cell migration upon interaction with PLD, FAK and Src. Journal of cell science 2015; 128(3): 516–526; e-pub ahead of print 2014/12/17; doi 10.1242/jcs.157339.

54. Fiume R, Ramazzotti G, Teti G, Chiarini F, Faenza I, Mazzotti G et al. Involvement of nuclear PLCbeta1 in lamin B1 phosphorylation and G2/M cell cycle progression. FASEB journal : official publication of the Federation of American Societies for Experimental Biology 2009; 23(3): 957–966; e-pub ahead of print 2008/11/26; doi 10.1096/fj.08-121244.

55. Ramazzotti G, Faenza I, Follo MY, Fiume R, Piazzi M, Giardino R et al. Nuclear phospholipase C in biological control and cancer. Crit Rev Eukaryot Gene Expr 2011; 21(3): 291–301; e-pub ahead of print 2011/11/25; doi 10.1615/critreveukargeneexpr.v21.i3.50.

56. Grewal S, Morrison EE, Ponnambalam S, Walker JH. Nuclear localisation of cytosolic phospholipase A_2_-α in the EA.hy.926 human endothelial cell line is proliferation dependent and modulated by phosphorylation. Journal of cell science 2002; 115(23): 4533; doi 10.1242/jcs.00146.

57. Hunt AN. Dynamic lipidomics of the nucleus. J Cell Biochem 2006; 97(2): 244-251; e-pub ahead of print 2005/10/22; doi 10.1002/jcb.20691.

58. Albi E, Viola Magni MP. The role of intranuclear lipids. Biol Cell 2004; 96(8): 657–667; e-pub ahead of print 2004/11/03; doi 10.1016/j.biolcel.2004.05.004.

59. Cascianelli G, Villani M, Tosti M, Marini F, Bartoccini E, Magni MV et al. Lipid microdomains in cell nucleus. Molecular biology of the cell 2008; 19(12): 5289–5295; e-pub ahead of print 2008/10/17; doi 10.1091/mbc.E08-05-0517.

60. Yoon MS, Rosenberger CL, Wu C, Truong N, Sweedler JV, Chen J. Rapid mitogenic regulation of the mTORC1 inhibitor, DEPTOR, by phosphatidic acid. Mol Cell 2015; 58(3): 549–556; e-pub ahead of print 2015/05/06; doi 10.1016/j.molcel.2015.03.028.

61. Panda A, Thakur R, Krishnan H, Naik A, Shinde D, Raghu P. Functional analysis of mammalian phospholipase D enzymes. Bioscience Reports 2018; 38(6); doi 10.1042/BSR20181690.

62. Gayral S, Deleris P, Laulagnier K, Laffargue M, Salles JP, Perret B et al. Selective activation of nuclear phospholipase D-1 by g protein-coupled receptor agonists in vascular smooth muscle cells. Circ Res 2006; 99(2): 132–139; e-pub ahead of print 2006/06/17; doi 10.1161/01.RES.0000232323.86227.8b.

63. Morris AJ. Phospholipases D: making sense of redundancy and duplication. Bioscience Reports 2019; 39(6); doi 10.1042/BSR20181883.

64. Barboro P, Ferrari N, Capaia M, Petretto A, Salvi S, Boccardo S et al. Expression of nuclear matrix proteins binding matrix attachment regions in prostate cancer. PARP-1: New player in tumor progression. International journal of cancer 2015; 137(7): 1574–1586; e-pub ahead of print 2015/03/27; doi 10.1002/ijc.29531.

65. Barboro P, Repaci E, D’Arrigo C, Balbi C. The role of nuclear matrix proteins binding to matrix attachment regions (Mars) in prostate cancer cell differentiation. PloS one 2012; 7(7): e40617; e-pub ahead of print 2012/07/19; doi 10.1371/journal.pone.0040617.

66. Ochs RL, Lischwe MA, Spohn WH, Busch H. Fibrillarin: a new protein of the nucleolus identified by autoimmune sera. Biol Cell 1985; 54(2): 123–133; e-pub ahead of print 1985/01/01; doi 10.1111/j.1768-322x.1985.tb00387.x.

67. Hui L, Abbas T, Pielak RM, Joseph T, Bargonetti J, Foster DA. Phospholipase D elevates the level of MDM2 and suppresses DNA damage-induced increases in p53. Molecular and cellular biology 2004; 24(13): 5677–5686; e-pub ahead of print 2004/06/17; doi 10.1128/mcb.24.13.5677-5686.2004.

68. Winter JN, Fox TE, Kester M, Jefferson LS, Kimball SR. Phosphatidic acid mediates activation of mTORC1 through the ERK signaling pathway. American journal of physiology Cell physiology 2010; 299(2): C335–344; doi 10.1152/ajpcell.00039.2010.

69. Rizzo MA, Shome K, Watkins SC, Romero G. The recruitment of Raf-1 to membranes is mediated by direct interaction with phosphatidic acid and is independent of association with Ras. The Journal of biological chemistry 2000; 275(31): 23911–23918; e-pub ahead of print 2000/05/10; doi 10.1074/jbc.M001553200.

70. Hein N, Hannan KM, George AJ, Sanij E, Hannan RD. The nucleolus: an emerging target for cancer therapy. Trends Mol Med 2013; 19(11): 643–654; e-pub ahead of print 2013/08/21; doi 10.1016/j.molmed.2013.07.005.

71. RamÍrez-Valle F, Badura ML, Braunstein S, Narasimhan M, Schneider RJ. Mitotic raptor promotes mTORC1 activity, G(2)/M cell cycle progression, and internal ribosome entry site-mediated mRNA translation. Molecular and cellular biology 2010; 30(13): 3151–3164; e-pub ahead of print 2010/05/05; doi 10.1128/mcb.00322-09.

72. Lyo D, Xu L, Foster DA. Phospholipase D stabilizes HDM2 through an mTORC2/SGK1 pathway. Biochemical and biophysical research communications 2010; 396(2): 562–565; e-pub ahead of print 2010/05/08; doi 10.1016/j.bbrc.2010.04.148.

73. Mathews TP, Hill S, Rose KL, Ivanova PT, Lindsley CW, Brown HA. Human phospholipase D activity transiently regulates pyrimidine biosynthesis in malignant gliomas. ACS chemical biology 2015; 10(5): 1258–1268; e-pub ahead of print 2015/02/04; doi 10.1021/cb500772c.

74. Liu Y, Kach A, Ziegler U, Ong AC, Wallace DP, Arcaro A et al. The role of phospholipase D in modulating the MTOR signaling pathway in polycystic kidney disease. PloS one 2013; 8(8): e73173; e-pub ahead of print 2013/09/07; doi 10.1371/journal.pone.0073173.

75. Hwang WC, Kim MK, Song JH, Choi K-Y, Min DS. Inhibition of phospholipase D2 induces autophagy in colorectal cancer cells. Experimental & Molecular Medicine 2014; 46(12): e124–e124; doi 10.1038/emm.2014.74.

76. Fiona M. Frame ARN, Sandra Klein, Rakesh Suman, Richard Kasprowicz, Vin M. Mann, Matt S. Simms, Norman J. Maitland. Tumor heterogeneity and therapy resistance - implications for future treatments of prostate cancer. Journal of Cancer Metastasis Treatment 2017; 3: 302–314; doi 10.20517/2394-4722.2017.34.

77. Loonen AJM, Soudijn W. Halopemide, a new psychotropic agent. Pharmaceutisch Weekblad (journal article) 1985; 7(1): 1–9; doi 10.1007/bf01962862.

78. Lindsley CW, Brown HA. Phospholipase D as a Therapeutic Target in Brain Disorders. Neuropsychopharmacology 2012; 37(1): 301–302.

