## Supplemental Figure 1 and legend for "Phospholipases D1 and D2 regulate cell cycling in primary prostate cancer cells and are differentially associated with the nuclear matrix": Bioarchive legend for supplementary Fig 1.docx

**Supplementary Figure 1**. Identification of PLD1 in PC3 cells at each stage of the nuclear matrix extraction procedure. PLD1 protein is detected in the nuclei of PC3 prostate cancer cells after detergent, high salt and DNase treatment but is lost after RNase digestion, red = PLD1, blue = DAPI to detect chromatin, green = lamin to define the perimeter of the nucleus. Scale bar 25mm. See Methodology for details.
