## Supplementary figures and images for "Phospholipases D1 and D2 regulate cell cycling in primary prostate cancer cells and are differentially associated with the nuclear matrix"

### Bioarchive supplementary Figure 1.pptx

#### Slide 1
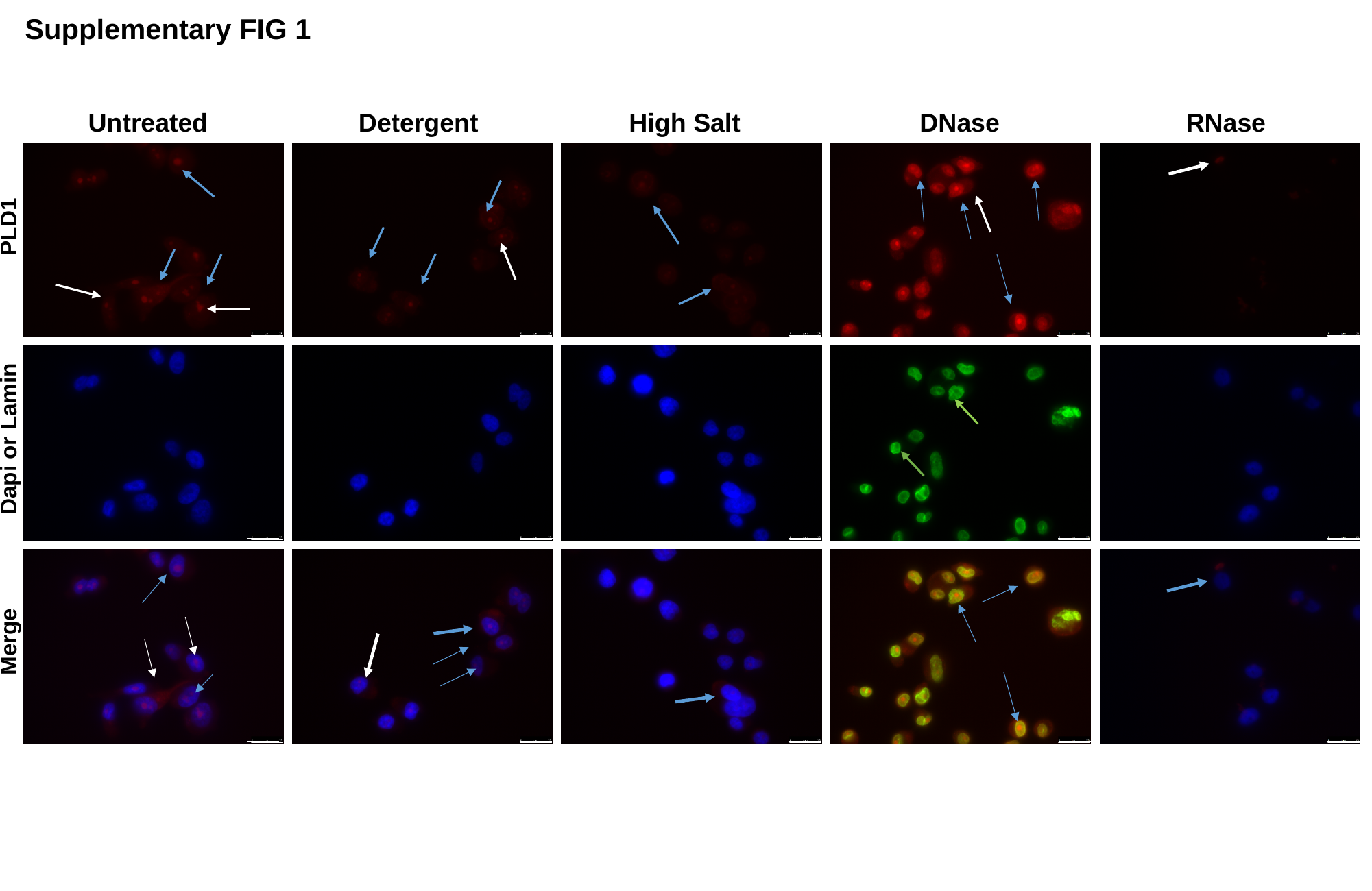

Supplementary FIG 1
 Untreated Detergent High Salt DNase RNase
PLD1
Dapi or Lamin
Merge
